# Targeting of Protein Kinase CK2 in Acute Myeloid Leukemia Cells Using the Clinical-Grade Synthetic-Peptide CIGB-300

**DOI:** 10.1101/2021.05.19.444866

**Authors:** Mauro Rosales, George V. Pérez, Ailyn C. Ramón, Yiliam Cruz, Arielis Rodríguez-Ulloa, Vladimir Besada, Yassel Ramos, Dania Vázquez-Blomquist, Evelin Caballero, Daylen Aguilar, Luis J. González, Katharina Zettl, Jacek R. Wiśniewski, Yang Ke, Yasser Perera, Silvio E. Perea

**Affiliations:** Department of Animal and Human Biology, Faculty of Biology, University of Havana (UH), Havana 10400, Cuba; Molecular Oncology Group, Department of Pharmaceuticals, Biomedical Research Division, Center for Genetic Engineering and Biotechnology (CIGB), Havana 10600, Cuba; Mass Spectrometry Laboratory, Proteomics Group, Department of Systems Biology, Biomedical Research Division, CIGB, Havana 10600, Cuba; Pharmacogenomic Group, Department of System Biology, Biomedical Research Division, CIGB, Havana 10600, Cuba; Biochemical Proteomics Group, Department of Proteomics and Signal Transduction, Max-Planck Institute of Biochemistry, Munich, Germany; China-Cuba Biotechnology Joint Innovation Center (CCBJIC), Yongzhou Zhong Gu Biotechnology Co., Ltd, Lengshuitan District, Yongzhou City 425000, Hunan Province, China

**Keywords:** protein kinase CK2, acute myeloid leukemia, CIGB-300, phosphoproteomics

## Abstract

Protein kinase CK2 has emerged as an attractive therapeutic target in acute myeloid leukemia (AML), advent that becomes particularly relevant since the treatment of this hematological neoplasia remains challenging. Here we explored for the first time the effect of the clinical-grade peptide-based CK2 inhibitor CIGB-300 on AML cells proliferation and viability. CIGB-300 internalization and subcellular distribution were also studied, and the role of B23/nucleophosmin 1 (NPM1), a major target for the peptide in solid tumors, was addressed by knock-down in model cell lines. Finally, pull-down experiments and phosphoproteomic analysis were performed to study CIGB-interacting proteins and identify the array of CK2 substrates differentially modulated after treatment with the peptide. Importantly, CIGB-300 elicited a potent anti-proliferative and proapoptotic effect in AML cells, with more than 80% of peptide transduced cells within three minutes. Unlike solid tumor cells, NPM1 did not appear to be a major target for CIGB-300 in AML cells. However, in vivo pull-down experiments and phosphoproteomic analysis evidenced that CIGB-300 targeted the CK2α catalytic subunit, different ribosomal proteins, and inhibited the phosphorylation of a common CK2 substrates array among both AML backgrounds. Remarkably, our results not only provide cellular and molecular insights unveiling the complexity of the CIGB-300 anti-leukemic effect in AML cells, but also reinforce the rationale behind the pharmacologic blockade of protein kinase CK2 for AML targeted therapy.

## 1. Introduction

Acute myeloid leukemia (AML) is a malignant disease characterized by infiltration of the blood, bone marrow, and other tissues by highly proliferative and abnormally differentiated myeloid progenitors [1, 2]. The origin of this disease has been associated with mutations affecting genes in different functional categories [3, 4]. For instance, mutations in genes encoding epigenetic modifiers connected to differentiation of myeloid progenitors are commonly acquired early, while mutations in protein kinases and other signaling molecules involved in cell proliferation and survival are typically secondary events [3, 4]. Regarding AML therapeutics, after no considerable changes in almost four decades, the treatment of patients with this hematological malignancy has recently seen some modifications with the approval for the FDA of several non-cytostatic agents [5]. This trend not only improves the therapeutic scenario for this neoplasia, but also highlights the suitability of molecular targeted drugs in AML [6]. Despite such rapid progress, the development of more efficient therapeutic approaches is still needed, mostly for AML patients with no actionable mutations or those with high risk of treatment-related side effect and mortality [7].

In such context, protein kinase CK2 has emerged as a valuable molecular target in the landscape of protein kinases with pivotal role in AML biology [8–11]. CK2 is a highly conserved and ubiquitously expressed serine/threonine protein kinase that regulates canonical cellular processes in cell physiology [12, 13]. In human cells, this enzyme exists as a tetrameric structure comprising two catalytic (α or its isoform α’) and two regulatory (β) subunits [14]. Phosphorylation by CK2 is specified by several acidic residues located downstream from the phosphoacceptor amino acid, the one at position n + 3 playing the most important function [15, 16]. Conversely, basic residues at any position close to phosphoacceptor amino acid and proline at position n + 1, exerts negative effect over CK2-mediated phosphorylation [17].

Protein kinase CK2 is responsible of roughly 25% of cellular phosphoproteome, and among its substrates are proteins implicated in transcription, translation, control of protein stability and degradation, cell cycle progression, cell survival, circadian rhythms, and virus replication [18]. Such extraordinary pleiotropy, explain why CK2 have been linked to multiple human diseases including neurological and psychiatric disorders, viral infections as well as solid and hematological neoplasia [19–21]. In fact, CK2 modulates a number of signaling pathways essential for hematopoietic cell survival and function, and high expression of CK2α subunit has been associated with lower disease-free survival and overall survival rates in AML patients with normal karyotype [22, 23]. Moreover, leukemic cells are significantly more sensitive to CK2 downregulation as demonstrated using genetic and pharmacologic approaches [8, 24]. The latter becomes particularly relevant, since myeloid malignant diseases stand among the most aggressive and lethal types of cancer, and are often characterized by resistance to standard chemotherapy as well as poor long-term outcomes [24].

In line with mounting evidences supporting the instrumental role of CK2 in human malignancies, various strategies to inhibit its activity have been explored in pre-clinical studies. However, only two compounds, the ATP-competitive inhibitor CX-4945 and the synthetic-peptide CIGB-300, have reached clinical trials [25, 26]. CX-4945 is a selective ATP-competitive CK2 inhibitor that has shown antineoplastic effect in solid tumor and hematological malignancies [24, 27]. This orally bioavailable small-molecule has been tested in Phase I clinical trial in patients with advanced solid tumors and Phase I/II randomized clinical trial in patients with cholangiocarcinoma [25, 28, 29]. On the other hand, CIGB-300 is a chimeric peptide containing a cell-penetrating moiety, that was originally designed to block the CK2-mediated phosphorylation through binding to phosphoacceptor domain in the substrates [26, 30]. Remarkably, the nucleolar protein B23/nucleophosmin 1 (NPM1) has been suggested as a major target for CIGB-300 in solid tumor [31, 32]. However, pull-down experiments and phosphoproteomics analysis have recently evidenced that the CIGB-300 mechanism could be more complex than originally thought [33, 34]. Such studies demonstrated that the peptide can interact with protein kinase CK2α catalytic subunit and impair CK2 enzymatic activity in non-small cell lung cancer (NSCLC) and T-cell acute lymphoblastic leukemia (T-ALL) cell lines [33, 34]. Concerning CIGB-300 antineoplastic effect, the peptide has exhibited a strong pro-apoptotic and anti-tumor effect in pre-clinical cancer models [35, 36], and has also been tested in Phase I/II clinical trial in patients with cervical cancer and Phase I trial in patients with relapsed/refractory solid tumors [37–40].

Regarding hematological neoplasia, the anti-leukemic effect of CK2 inhibitor CX-4945 has been evaluated in several studies comprising both lymphoid and myeloid malignancies [11, 24, 41–43], whereas CIGB-300 has only been explored in chronic and acute lymphocytic leukemia [34, 44]. Therefore, we decided to evaluate the potential anti-neoplastic effect of CIGB-300 in AML cells. Interestingly, we demonstrated that CIGB-300 exerts a potent antiproliferative and pro-apoptotic effect in AML cells, and provided a group of cellular and molecular evidences supporting the anti-leukemic effect of the peptide in this hematological malignancy.

## 2. Materials and Methods

### 2.1. Cell Culture

Human AML cell lines HL-60 and THP-1 were originally obtained from the American Type Culture Collection (ATCC, Manassas, VA, USA), while OCI-AML3, EOL-1 and K562 were obtained from the German Collection of Microorganisms and Cell Cultures (DSMZ, Braunschweig, Germany). All cell lines were cultured in RPMI 1640 medium (Invitrogen, Carlsbad, CA, USA) supplemented with 10% (v/v) fetal bovine serum (FBS, Invitrogen, Carlsbad, CA, USA) and 50 μg/mL gentamicin (Sigma, St. Louis, MO, USA). All cells were maintained at 37 °C and 5% CO_2_.

### 2.2. Patient Samples

Bone marrow samples from five AML patients with high leukemia involvement were collected in accordance with the Declaration of Helsinki after informed consent and ethical approval of the Clinical Research Board from the Center for Genetic Engineering and Biotechnology (CIGB, Havana, Cuba) (Table S1). Mononuclear cells from bone marrow samples were isolated using Ficoll-Paque (GE Healthcare, Chicago, IL, USA) density gradient centrifugation. Cells were maintained under standard cell culture conditions.

### 2.3. AlamarBlue Assay

Proliferation of AML cell lines was determined using alamarBlue assay (Life Technologies, Carlsbad, CA, USA). Cells were seeded in flat-bottom 96-well plates (2 × 10^5^ cells/mL, 200 μL/well) and 24 h later serial dilutions 1:2 ranging from 100 to 3.12 μM of CIGB-300 were added. After 48 h of incubation, alamarBlue was added at 10% (v/v), and cell suspension were further incubated for 4 h. Fluorescence was measured in a CLARIOstar microplate reader (BMG LABTECH, Ortenberg, Germany) and half-inhibitory concentrations (IC_50_) values were estimated using CalcuSyn software (v2.1) (Biosoft, Cambridge, United Kingdom).

### 2.4. Annexin V/PI Staining

Viability of AML cells was measured using FITC Annexin V Apoptosis Detection Kit I (BD Biosciences, San Jose, CA, USA). Briefly, cell lines were incubated with 40 μM CIGB-300 for 3 and 5 h, while primary cells were incubated during 48 h. Following incubation, cells were washed twice with cold PBS and resuspended in binding buffer (1×) at a final concentration of 1 × 10^6^ cells/mL. Subsequently, 5 μL of FITC Annexin V and PI were added and cells suspensions were incubated for 15 min at room temperature in the dark. Flow cytometric analysis of stained cells was performed in Partec CyFlow Space instrument (Sysmex Partec GmbH, Gorlitz, Germany) and FlowJo software (v7.6.1) (BD, Ashland, OR, USA) was used for data analysis and visualization.

### 2.5. Cell Cycle Analysis

For cell cycle analysis, AML cells were incubated with 40 μM CIGB-300 during 5 and 24 h. Following treatment with the peptide, cells were collected by centrifugation, washed with PBS and fixed at 4 °C for 30 min with ice-cold 70% ethanol. After fixation, cells were treated with DNase-free RNase A (Sigma, St. Louis, MO, USA) and subsequently stained at 37 °C for 20 min with 50 μg/mL PI solution (Sigma, St. Louis, MO, USA). Stained cells were analyzed in the abovementioned Partec CyFlow Space flow cytometry and FlowJo Software (v7.6.1) was used for data processing and visualization.

### 2.6. Peptide Internalization and Confocal Microscopy

The internalization of CIGB-300 in AML cells was studied at 3, 10, 30 and 60 min after addition of 30 μM of the peptide with N-terminal fluorescein tag (CIGB-300-F). Once incubated with the peptide, cells were washed and resuspended in PBS solution containing 50 μg/mL PI (Sigma, St. Louis, MO, USA), to discriminate viable cells from cells with membrane damage. Finally, flow cytometric analysis was performed as previously described. Alternatively, to study the peptide subcellular distribution, cells were seeded in 6-well plates and 24 h later CIGB-300-F and Hoechst 33342 (Sigma, St. Louis, MO, USA) were added at a final concentration of 30 μM and 4 μg/mL, respectively. After 10, 30 and 60 min of incubation under standard culture conditions, living cells were examined under Olympus FV1000 confocal laser scanning microscope (Olympus, Tokyo, Japan). Images were acquired with UPLSAPO 60× immersion objective and processed using Olympus FluoView software (v4.0) (Olympus, Tokyo, Japan).

### 2.7. Lentiviral Infection

Cells were infected with HIV-based third-generation lentiviral particles according to experimental conditions previously described [34]. Briefly, a shRNA sequence against the 3’UTR region of *NPM1* was inserted into the lentiviral transfer plasmid pLG. This plasmid contains a GFP reporter gene and lentiviral vector elements to produce infective particles when co-transfected with packaging plasmids pLP1, pLP2 and pLP/VSVG in HEK-293T cells (ViraPower Lentiviral Packaging Mix, Thermo Fisher Scientific, Waltham, MA, USA). After virus production and subsequent titration, cells were infected by spinoculation in the presence of polybrene (8 μg/mL) and at 10× multiplicity of infection [45]. On the next day, fresh medium was added and the cells were allowed to recover for another 48 h. Finally, infected cells were analyzed by flow cytometry and western blot for evaluation of GFP expression and NPM1 protein levels, respectively.

### 2.8. Pull-Down Experiments

For in vivo pull-down assays, CIGB-300 with N-terminal biotin tag (CIGB-300-B) was added to AML cells at a final concentration of 40 μM and incubated for 30 min. Subsequently, cells were collected, washed and lysed in hypotonic PBS solution (0.1×) containing 1 mM DTT (Sigma, St. Louis, MO, USA) and complete protease inhibitor (Roche, Basel, Switzerland) by eight freeze-thaw (liquid nitrogen/37 °C) cycles. Cellular lysates were cleared by centrifugation and 300 μg of total protein were added to 30 μL of pre-equilibrated streptavidin-sepharose matrix (GE Healthcare, Chicago, IL, USA). Following 1 h at 4 °C, the matrix was collected by centrifugation and extensively washed with PBS solution (1×) containing 1 mM DTT. Proteins bound to streptavidin-sepharose matrix were digested with trypsin (Promega, Madison, WI, USA) during 16 h or eluted for western blot analysis. In parallel, untreated cells were subjected to the same experimental procedure to identify those proteins non-specifically bound to streptavidin-sepharose matrix.

### 2.9. Western Blot

Cells were lysed in RIPA buffer containing protease/phosphatase inhibitor (Thermo Fisher Scientific, Waltham, MA, USA), and equal amounts of protein were resolved in 12.5% SDS-PAGE [46]. Next, proteins were transferred to a nitrocellulose membrane and immunoblotted with the following antibodies according to instructions from the manufacturer: NPM1, p-NPM1 (S125) and CK2α (Abcam, Cambridge, United Kingdom), β-actin (Sigma, St. Louis, MO, USA). Detection was performed with peroxidase-conjugated anti-mouse or anti-rabbit IgG (Sigma, St. Louis, MO, USA), and signal was developed using SuperSignal West Pico Chemiluminescent Substrate (Thermo Fisher Scientific, Waltham, MA, USA).

### 2.10. Protein Identification by LC-MS/MS

After tryptic digestion, resulting peptides from treated and untreated samples were isotopically labelled with N-acetoxy-D_3_-succinimide (D_3_-NAS) and N-acetoxy-H_3_-succinimide (H_3_-NAS), respectively, and then pooled together. Finally, proteins were identified by LC-MS/MS analysis using Eksigent nanoLC (AB SCIEX, Framingham, MA, USA) coupled to LTQ Orbitrap Velos Pro mass spectrometer (Thermo Fisher Scientific, Waltham, MA, USA). Protein-protein interaction networks were constructed using information from STRING database [47], and protein kinase CK2 substrates were identified using post-translational modification resource iPTMnet, web-based tool KEA2 and literature search [48, 49].

### 2.11. Phosphoproteomic Analysis

Sample preparation, phosphopeptide enrichment and nanoLC-MS/MS for phosphoproteomic analysis of AML cells treated or not with 40 μM CIGB-300 during 30 min were conducted as previously described [50]. Briefly, three replicates from each condition were processed by multienzyme digestion filter-aided sample preparation (MED-FASP) with overnight lys-C and tryptic digestions [51], and the phosphopeptides enriched by TiO_2_ chromatography. Phosphopeptides were later injected through an EASY-nLC 1200 system into a QExactive HF mass spectrometer (Thermo Scientific, USA) using a home-made column (75 mm ID, 20 cm length), and separated with a gradient from 5% buffer B (0.1% formic acid in acetonitrile) up to 30% in 45 min, 30-60% in 5 min, and 60-95% in 5 min more.

### 2.12. Data Processing and Bioinformatics

Data processing and quantification were performed in MaxQuant software (v1.6.2.10) [52] and Perseus computational platform (v1.6.2.2) [53]. Phosphopeptides dataset were filtered of reverse and potential contaminants hits, and only phosphosites with localization probability above 0.75 were retained for further analysis. Besides, only those phosphopeptides represented in each group were considered, and differences in phosphosites occupancy higher than 25% were selected as significant changes. Enzyme-substrate relations were retrieved using post-translational modification resource iPTMnet and web-based tool KEA2 [48, 49]. In addition to well-documented substrates, we searched for candidate CK2 substrates based on the presence of the CK2 consensus sequence [16], the enzyme-substrate predictions retrieved from NetworKIN database [54], the dataset of high confidence CK2 substrates reported by Bian et al. [55], and the phosphoproteins that interact with protein kinase CK2 according to Metascape database [56].

### 2.13. Statistical Analysis

Differences between groups were determined using Student’s t Test or one-way ANOVA followed by Tukey’s multiple comparisons test. Data were analyzed and represented in GraphPad Prism (v6.01) (GraphPad Software, Inc, San Diego, CA, USA).

## 3. Results

### 3.1. Inhibition of CK2 Impairs AML Cells Proliferation and Viability

In order to assess the impact of CK2 inhibition in AML cells, we evaluated the effect of CIGB-300 peptide on proliferation and viability of five human cells lines representing multiple stages of myeloid differentiation and AML subtypes (**Figure 1**). As measured using alamarBlue assay, CIGB-300 exhibited a strong dose-dependent inhibitory effect on proliferation of AML cell lines, with IC_50_ values ranging from 21 to 33 μM (**Figure 1A**). Such antiproliferative effect could be explained in part by an impairment of AML cells viability, since Annexin V-FITC/PI staining indicated that the peptide promoted apoptosis in HL-60 and OCI-AML3 cells, and primary cells from the majority of AML patients bone marrow samples (**Figure 1B, C**). We also determined the impact CIGB-300 peptide on progression of AML cells through cell cycle. Our results evidenced that the peptide induced an accumulation of HL-60 cells in S phase, while no change in cell cycle distribution was detected in OCI-AML3 cells (**Figure 1D**).

**Figure 1.**
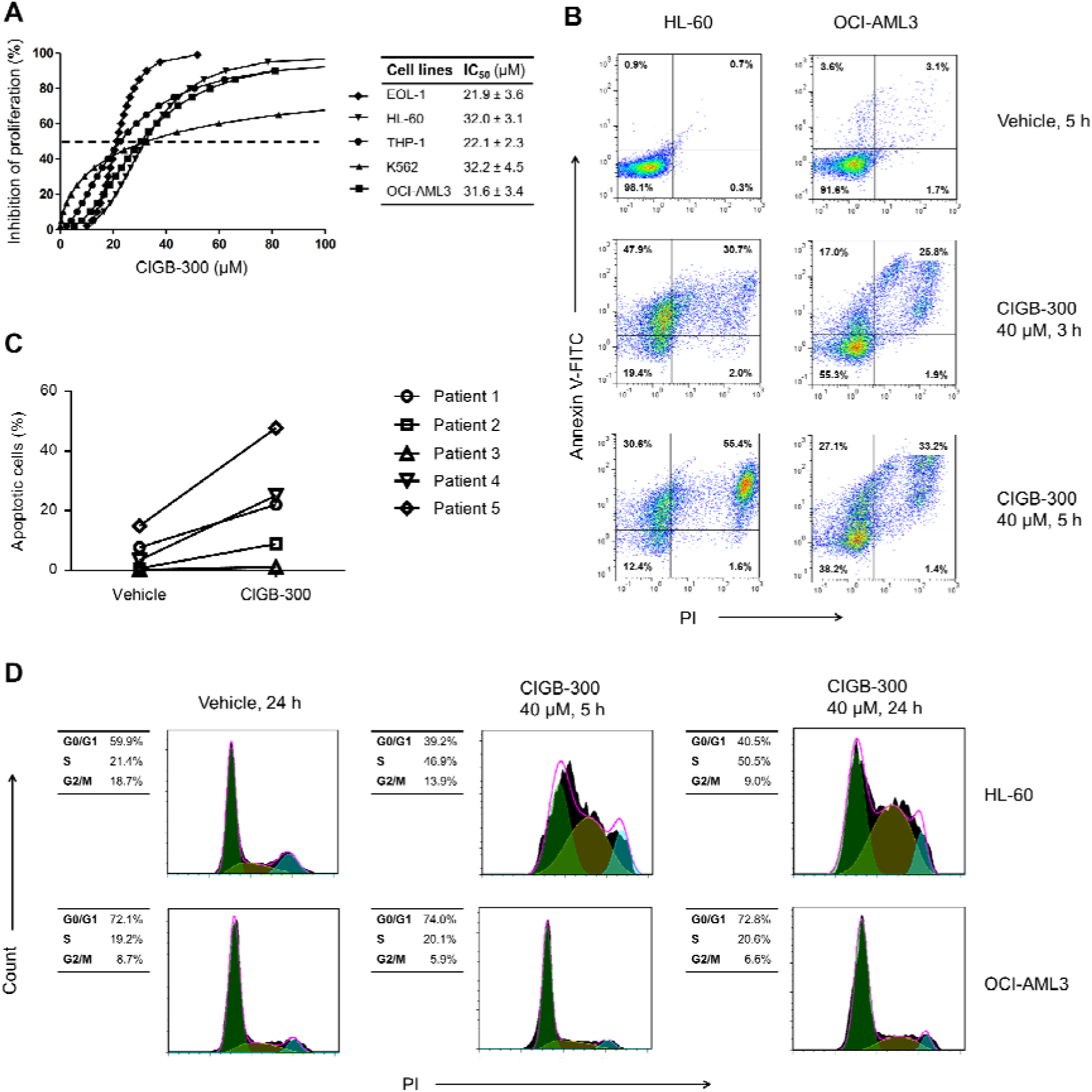
CIGB-300 peptide impairs proliferation and viability of AML cells: (**a**) Evaluation of CIGB-300 anti-proliferative effect in human cell lines representing different AML subtypes using alamarBlue assay; Viability of HL-60, OCI-AML3 cells (**b**) and primary cells from AML patients (**c**) was measured by Annexin V/PI staining after incubation with 40 μM CIGB-300 at the indicated time points; (**d**) Cell cycle analysis was conducted through PI staining followed by flow cytometry at 5 and 24 h of incubation with 40 μM CIGB-300. Results from (**a**) are shown as mean ± SD, n = 3.

### 3.2. Internalization and Subcellular Distribution of CIGB-300

To study CIGB-300 internalization and subcellular distribution, we conducted flow cytometry and confocal microscopy experiments using CIGB-300-F (**Figure 2**, **Figure S1**). We found that the peptide was rapidly internalized (within only 3 min) in more than 80% of HL-60 and OCI-AML3 cells (**Figure 2A**). Remarkably, the percentage of fluorescent cells maintained unchanged during the remaining incubation times (up to 60 min) (**Figure 2A**). Moreover, intracellular accumulation of CIGB-300 was determined in the fluorescein-positive population. As indicated the geometric mean (Gmean) of fluorescence intensity, after 3 and 10 min of incubation HL-60 cells showed higher intracellular levels of CIGB-300-F when compared to OCI-AML3 cells (**Figure 2B**), however, such difference disappeared after 30 and 60 min of incubation (**Figure 2B**).

**Figure 2.**
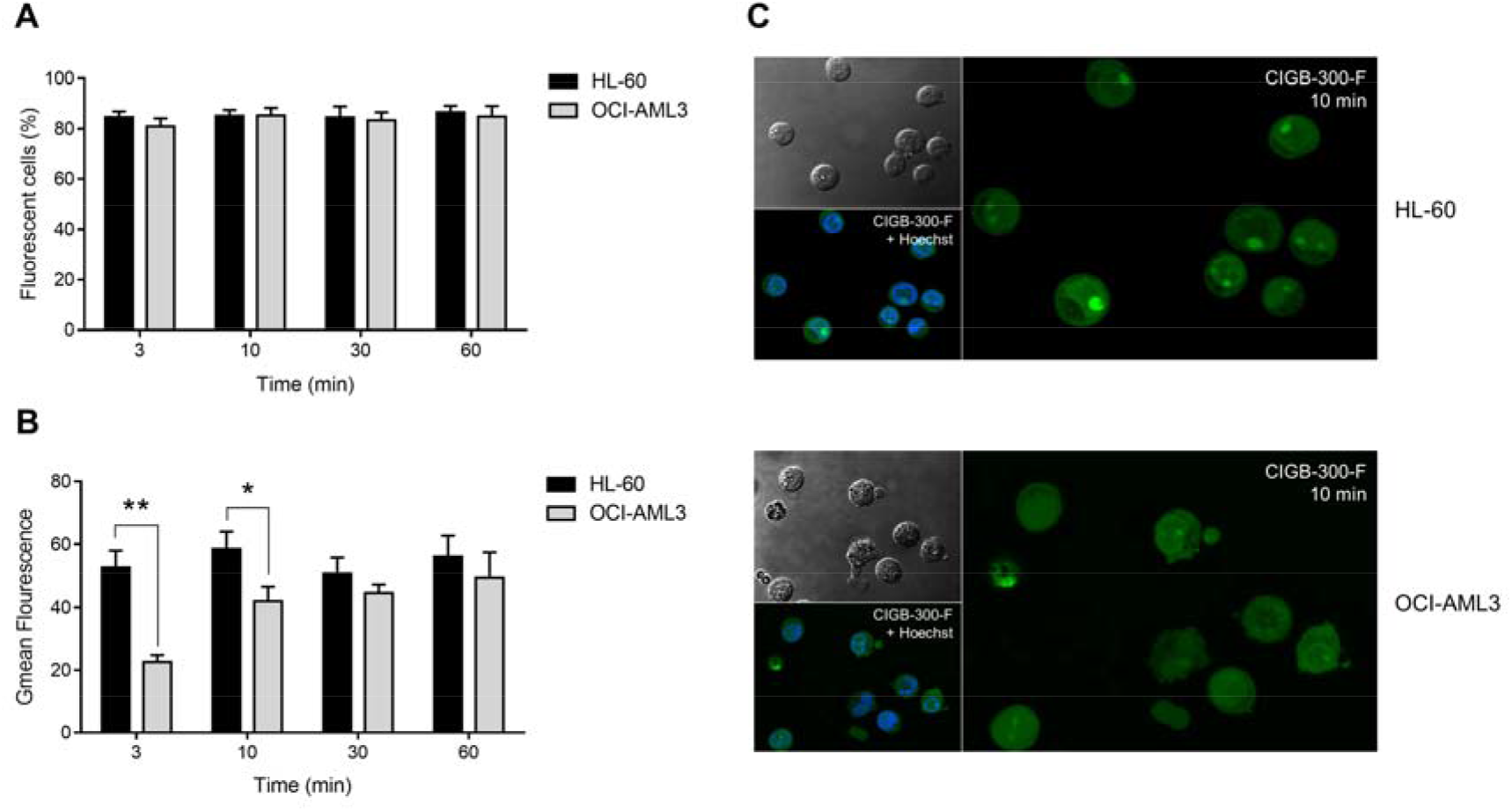
Internalization and subcellular distribution of CIGB-300 in AML cells: (**a**) Internalization of CIGB-300 in HL-60 and OCI-AML3 cells was studied using CIGB-300-F. Percentage of fluorescent cells was determined by flow cytometry after 3, 10, 30 and 60 min of incubation with 30 μM of CIGB-300-F; (**b**) Intracellular accumulation of CIGB-300 was measured as Gmean of fluorescence intensity in the fluorescein-positive population; (**c**) Subcellular distribution of CIGB-300 in AML cells after 10 min of incubation with 30 μM of CIGB-300-F. A total of 5 optical fields were examined for each experimental condition in confocal microscopy experiment. Results from (**a**) and (**b**) are shown as mean ± SD, n = 3. (*) *p*-value < 0.05, (**) *p*-value < 0.01.

Subcellular distribution of CIGB-300-F was also examined by confocal microscopy (**Figure 2C**, **Figure S1**). We found that in HL-60 cells the peptide preferentially accumulated in the nucleolus after 10 min, whereas the cytoplasm and the nucleoplasm displayed comparable fluorescence intensities (**Figure 2C**). In contrast, OCI-AML3 cells exhibited a more diffuse distribution pattern between the cytoplasm, the nucleoplasm and the nucleolus, with no significant accumulation at any of these subcellular locations (**Figure 2C**). Similar to transduction levels, subcellular distribution of CIGB-300-F was identical at the other assessed time points (30 and 60 min) (**Figure S1**).

### 3.3. Profiling CIGB-300 Interactome in AML Cells

Once demonstrated that CIGB-300 is readily internalized in AML cells, we explored the peptide interaction profile using in vivo pull-down experiments followed by LC-MS/MS analysis. Using this experimental approach, we identified a group of 48 and 70 proteins conforming the CIGB-300 interactome in HL-60 and OCI-AML3 cells, respectively, with an overlap of 40 proteins that were identified in both cellular backgrounds (**Figure 3**, **Table S2**). For a better understanding of CIGB-300 interactome we constructed PPI networks with identified proteins using information annotated in STRING database (**Figure 3**) [47]. In PPI network associated to HL-60 cells we detected one functional complex corresponding to the ribosome (37 structural proteins from the small and the large ribosome subunits); whereas in OCI-AML3, besides the ribosome (51 structural proteins), two more functional complexes corresponding to the nucleosome (4 proteins) and the spliceosome (6 proteins) appeared represented (**Figure 3**).

**Figure 3.**
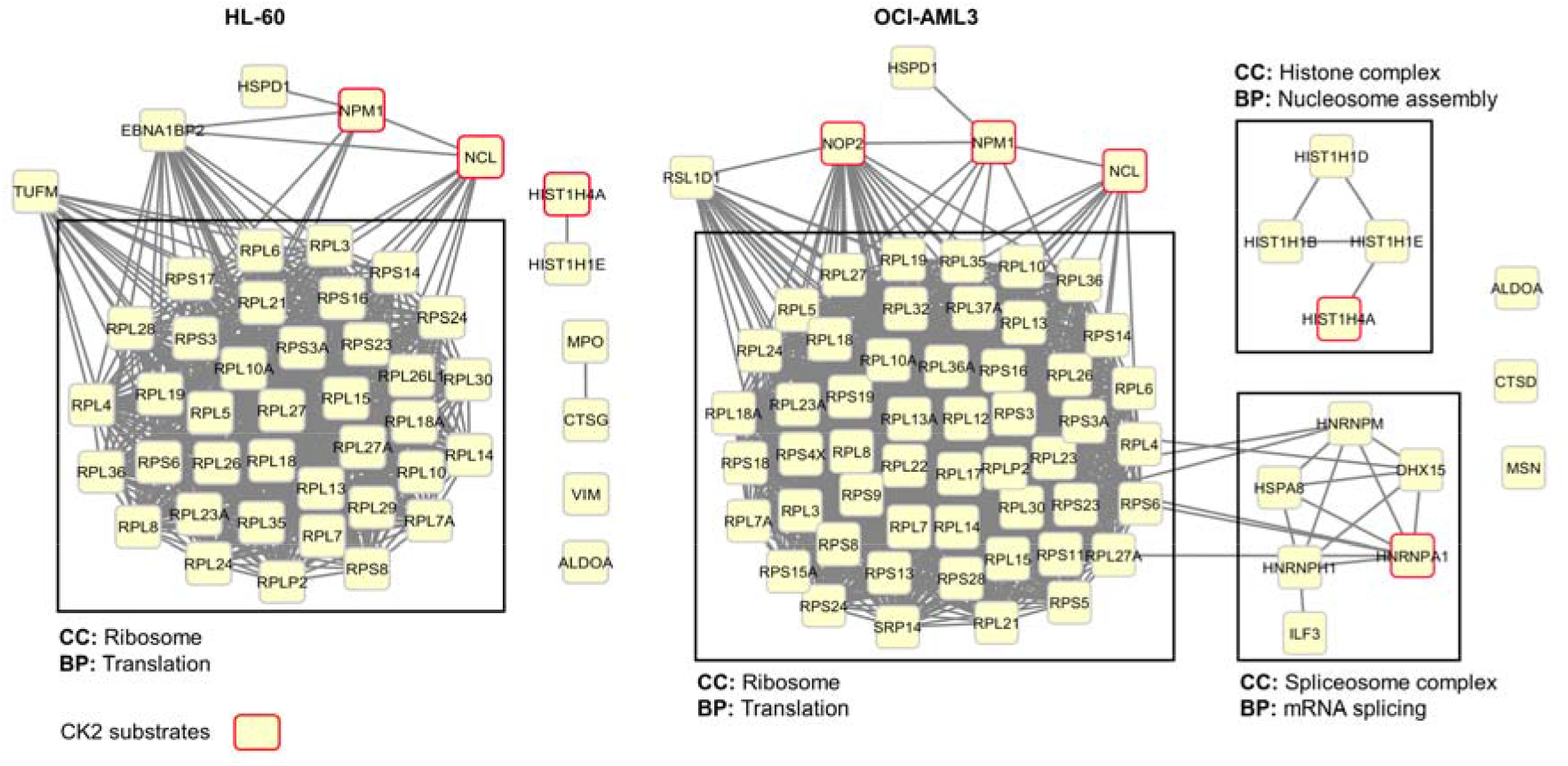
Protein-protein interaction networks associated to CIGB-300 interacting proteins in HL-60 and OCI-AML3 cells. Networks were generated using information gathered from STRING database and visualized using Cytoscape. Cellular components (CC) and biological processes (BP) retrieved from Gene Ontology database are indicated.

Among the subset of CIGB-300 interacting proteins, we searched for protein kinase CK2 substrates. In HL-60 cells the peptide interacted with three well-documented CK2 substrates according to iPTMnet and KEA databases and literature search, while in OCI-AML3 cells five substrates were identified (**Figure 3**, **Table S2**). Of note, the three CK2 substrates identified in HL-60 cells, nucleolar proteins NPM1 and nucleolin (NCL), as well as histone H4 (HIST1H4A), were also identified as part of CIGB-300 interactome in OCI-AML3 cells (**Figure 3**, **Table S2**).

### 3.4. NPM1 is Not a Major Target for CIGB-300 in AML Cells

Considering that NPM1 has been suggested as a major target for CIGB-300 in solid tumors [31, 32], we carried out phosphorylation experiments to corroborate the inhibition of NPM1 phosphorylation in the presence of CIGB-300. As expected, in both AML cell lines we detected the inhibition of CK2-mediated phosphorylation at S125 residue of NPM1 by western blot (**Figure S2**). To clarify the relevance of such inhibition, we further infected AML cells with lentiviral particles expressing a shRNA against the 3’-UTR of NPM1 mRNA (**Figure 4**). Once transduced, cells were analyzed by flow cytometry in order to determine the percent of GFP-positive (GFP+) cells as indicator of lentiviral infection efficiency (**Figure 4A**).

**Figure 4.**
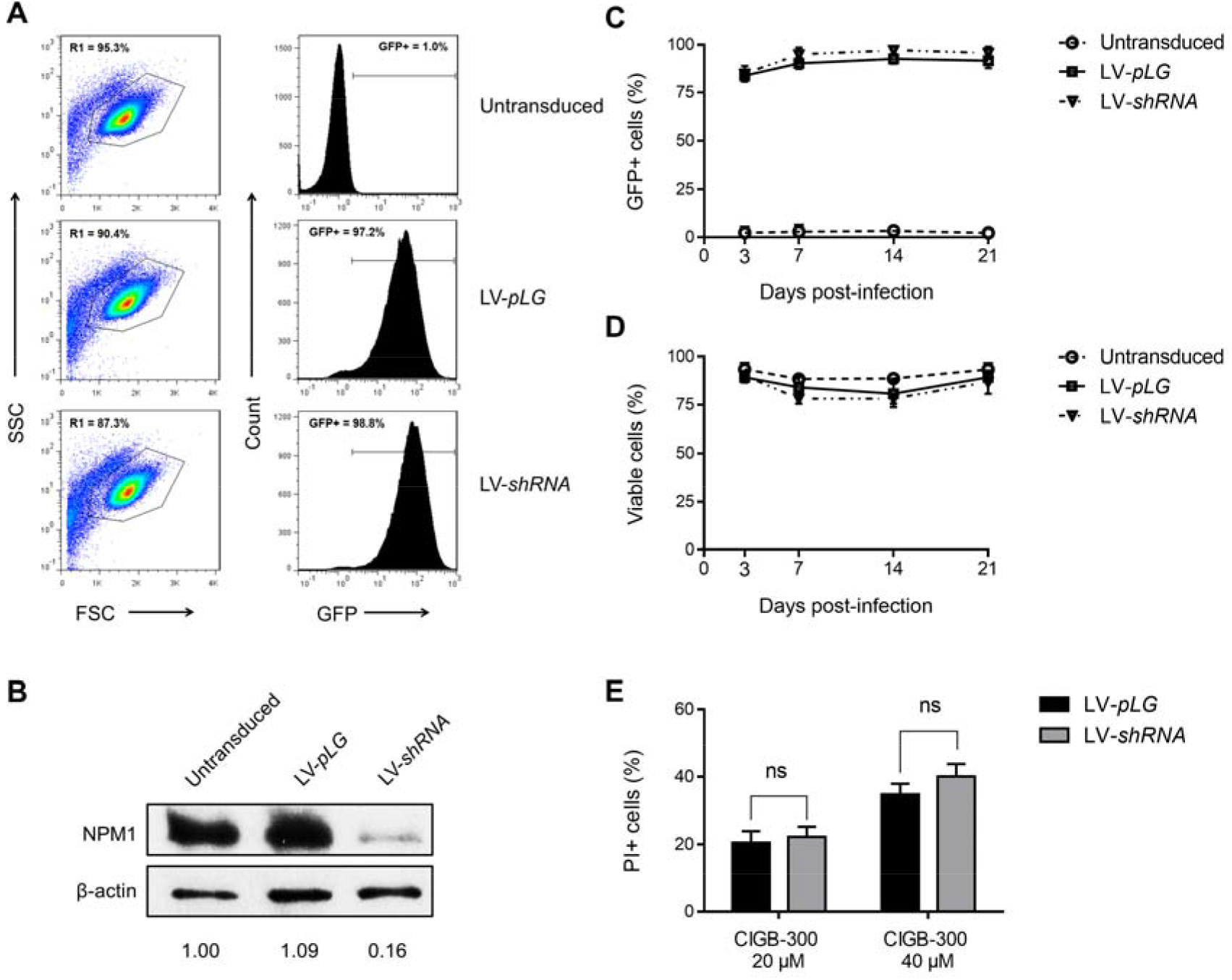
Lentiviral vector-mediated knock-down of NPM1 in AML cells: (**a**) Flow cytometry analysis of GFP expression in transduced HL-60 cells at day 7 post-infection. Percentage of GFP+ cells was determined on live-cell population (R1 gate in dot plots); (**b**) Immunoblots from HL-60 transduced cells showing NPM1 protein knock-down in LV-*shRNA* infected cells. Densitometry analysis values of NPM1 bands were normalized to β-actin and then to untransduced cells; (**c**) GFP expression levels and (**d**) viability of transduced HL-60 cells were followed by flow cytometry during three weeks post-infection; (**e**) Sensibility of transduced cells toward the cytotoxic effect of CIGB-300 was evaluated by PI staining. Cells were incubated with 20 or 40 μM of CIGB-300 for 5 h, stained with PI solution and then analyzed by flow cytometry. Results from (**c**), (**d**) and (**e**) are shown as mean ± SD, n = 3. (ns) not significant.

In accordance with infection efficiencies, western blot analysis evidenced a clear down-regulation of NPM1 protein levels (**Figure 4B**). HL-60 cells transduced with either empty vector (LV-*pLG*) or NPM1 silencing vector (LV-*shRNA*), showed similar infection efficiencies (higher than 95%), and consistently presented no differences in GFP expression or viability during at least three weeks post-infection (**Figure 4C, D**). Worthy of note that despite roughly 85% knock-down of NPM1 protein, HL-60 cells transduced with either LV-*pLG* or LV-*shRNA* had no differential sensibility toward the CIGB-300 cytotoxic effect (**Figure 4E**).

Finally, while in HL-60 cells more than 95% infection efficiency was achieved, we were unable to transduce OCI-AML3 cells. In view of such inconvenience, we conducted similar knock-down experiments using AML cell line THP-1, obtaining results similar to those described for HL-60 cells (**Figure S3**).

### 3.5. CIGB-300 Regulates the CK2-dependant Phosphoproteome

Given that NPM1 appears not to be a critical target for CIGB-300 in AML cells, and considering that direct enzyme inhibition has been pointed as a parallel mechanism for the peptide in NSCLC and T-ALL cells [33, 34], we carried out pull-down experiments followed by immunodetection of CK2α catalytic subunit. As recently described for other cancer cell lines, a clear interaction between CIGB-300 and CK2α subunit was detected in AML cells (**Figure 5A**). The aforementioned lead us to conduct phosphoproteomics analysis of HL-60 and OCI-AML3 cells treated with the peptide, in order to fully gauge the complexity of CIGB-300 inhibitory mechanism in AML cells (**Figure 5B**). As a result, in HL-60 cells we identified 5216 phosphopeptides corresponding to 4714 unique phosphorylation sites on 2267 proteins. On the other hand, in OCI-AML3 cells 3053 phosphopeptides corresponding to 2812 unique phosphorylation sites on 1531 proteins were identified. Overall, we identified 5460 unique phosphorylation sites corresponding to 2576 proteins, with an overlap of 2066 phosphorylation sites that were identified in both cell lines (**Figure 5C**, **Table S3**).

**Figure 5.**
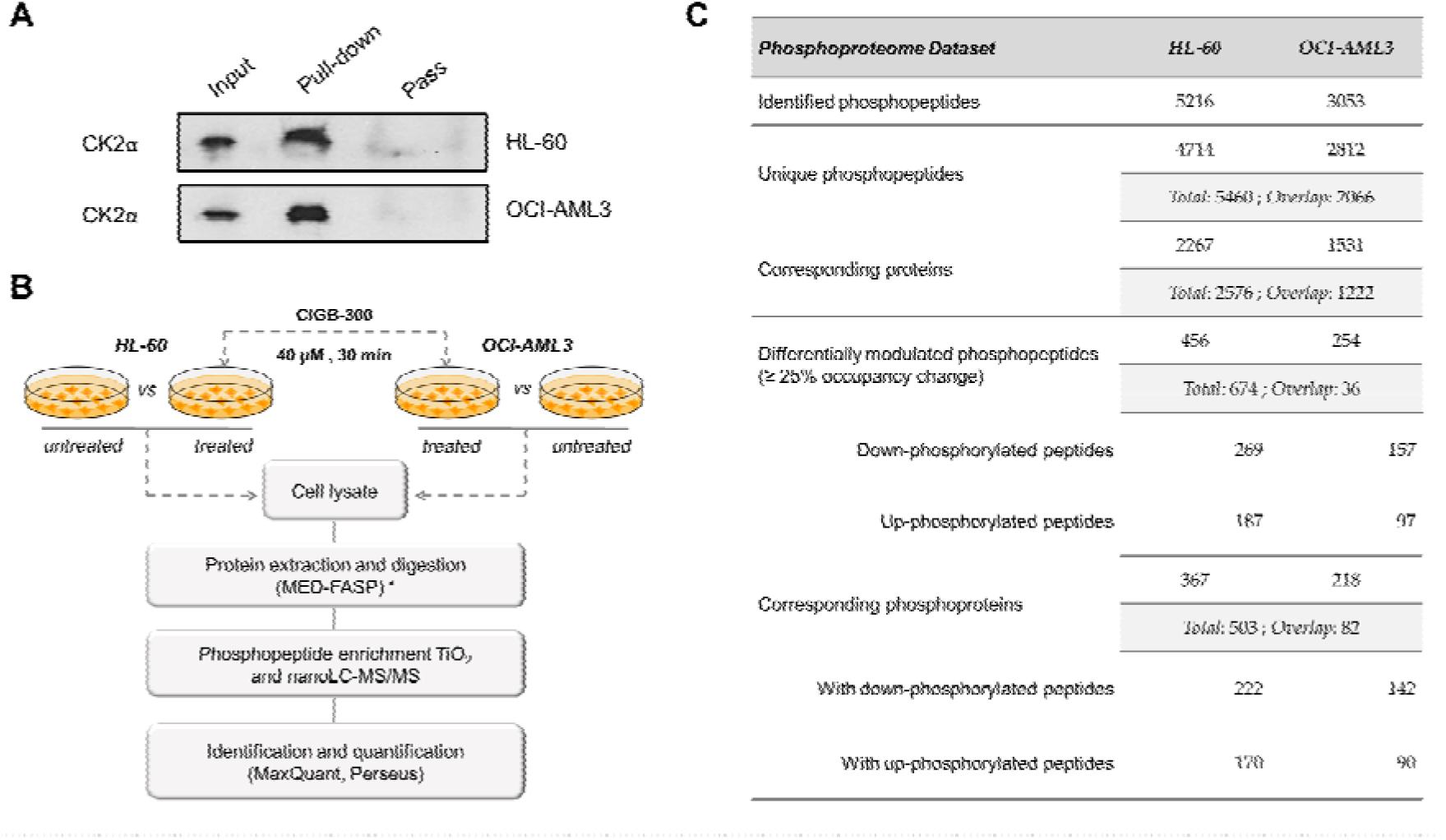
Phosphoproteomic analysis of human AML cells treated with the CK2 inhibitor CIGB-300: (**A**) Immunoblots from in vivo pull-down fractions to detect the interaction between CIGB-300 and CK2α catalytic subunit; (**B**) Workflow for the exploration of phosphorylation changes induced in HL-60 and OCI-AML3 cells after treatment with CIGB-300. Three biological replicates of each group were evaluated; (**C**) Number of identified and differentially modulated phosphopeptides in each AML cell line. (*) MED-FASP: multienzyme digestion filter-aided sample preparation [51]. Input: cellular extract, Pull-down: bound fraction, Pass: flow-through fraction.

In HL-60 cells, treatment with CIGB-300 significantly decreased phosphorylation site occupancy (25% or more) of 269 phosphopeptides belonging to 222 proteins, whereas 187 phosphopeptides belonging to 170 proteins showed increased occupancy (25% or more) (**Figure 5C**, **Table S4**). Similarly, in OCI-AML3 cells 157 phosphopeptides on 142 proteins underwent 25% or more phospho-occupancy reduction, while only 97 phosphopeptides on 90 proteins increased their occupancy (**Figure 5C**, **Table S4**). Among the differentially modulated phosphosites, 11 (7 down-phosphorylated) and 8 (3 down-phosphorylated) are reported as bona fide CK2 substrates in HL-60 and OCI-AML3 cells, respectively; and septin-2 phospho-serine 218 (SEPTIN2 S218), a well-documented CK2 substrate, appeared inhibited in both AML cell lines (**Table S5**). A significant number of phosphosites attributed to glycogen synthase kinase-3 beta (GSK3B) and members of mitogen-activated protein kinases (MAPKs) and cyclin-dependent kinases (CDKs) families, also appeared modulated in both phosphoproteomic profiles (**Table S5**).

In addition to substrates annotated in iPTMnet and KEA databases, we searched for candidate CK2 substrates among down-regulated phosphopeptides based on: 1) the presence of CK2 consensus sequence, 2) the enzyme-substrate predictions retrieved from NetworKIN database, 3) the dataset of high confidence CK2 substrates reported by Bian et al. [55], and 4) the phosphoproteins that interact with protein kinase CK2 according to Metascape database (**Table S6**). For instance, 63 and 31 phosphopeptides showing 25% or more occupancy reduction upon treatment with CIGB-300, fulfilled the protein kinase CK2 consensus sequence in HL-60 and OCI-AML3 cells, respectively (**Table S6**). Besides, 9 and 6 phosphopeptides from HL-60 and OCI-AML3 phosphoproteomic profiles were identified as CK2 substrates based on predictions retrieved from NetworKIN database. Finally, candidate CK2 substrates dataset was filtered out to find those substrates that had the concomitant occurrence of two or more criteria associated to CK2 phosphorylation (**Table S6**). Using such workflow, in HL-60 cells 34 phosphosites on 30 proteins were identified as the most reliable CK2 substrates modulated after treatment with CIGB-300, while 16 phosphosites on 14 proteins were identified in OCI-AML3 (**Table 1**, **Table S6**). The list also includes those CK2 substrates previously confirmed as bona fide according to iPTMnet and KEA databases.

**Table 1.**
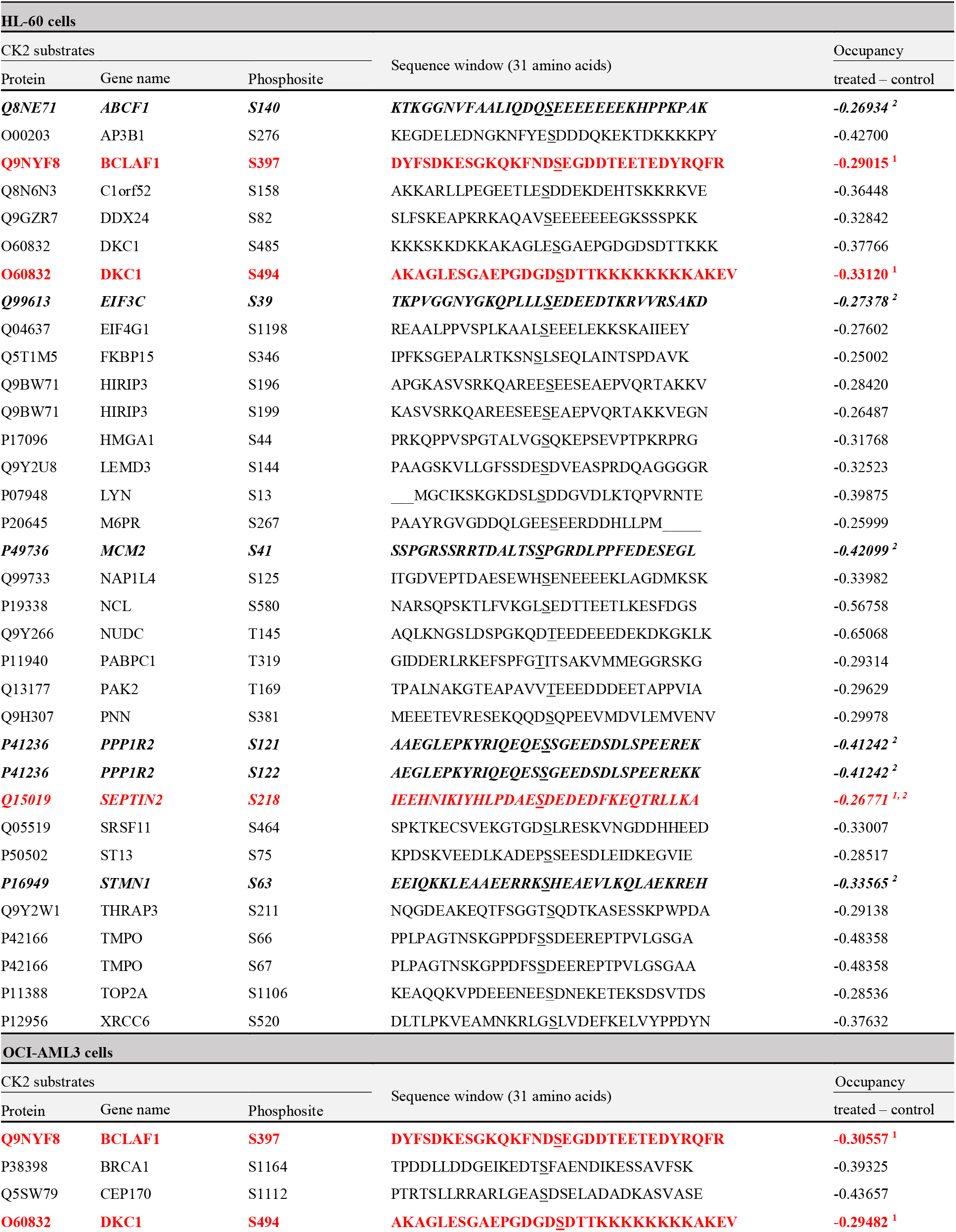
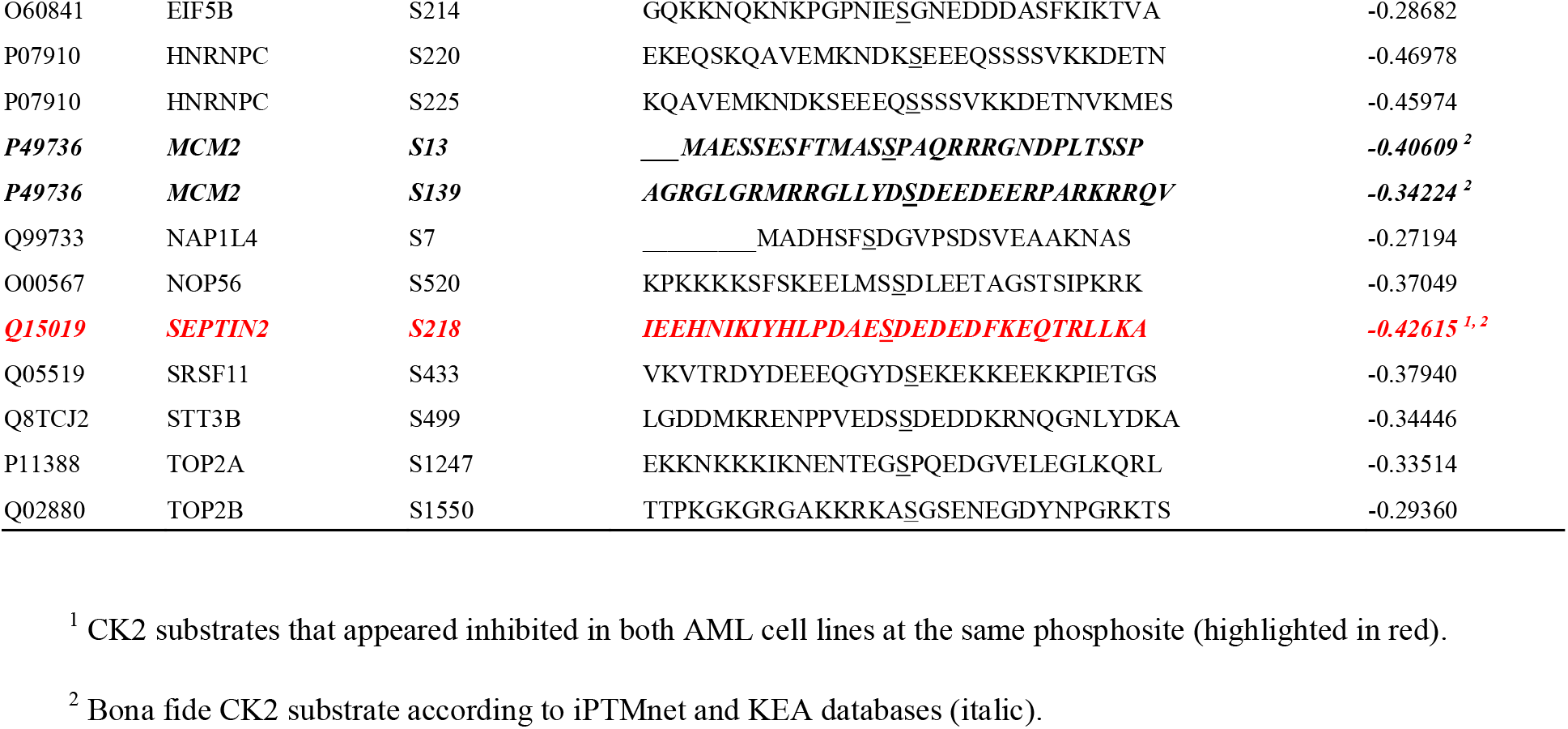
Summary of the most reliable CK2 substrates differentially modulated after treatment of AML cells with CIGB-300.

Importantly, besides the bona fide CK2 substrate SEPTIN S218, other candidate substrates considered among the most reliable such as Bcl-2-associated transcription factor 1 S397 (BCLAF1 S397) and H/ACA ribonucleoprotein complex subunit 4 S494 (DKC1 S494), were identified in HL-60 and OCI-AML3 cells with similar phospho-occupancy reduction (**Table 1**, **Table S6**). Furthermore, DNA replication licensing factor MCM2, nucleosome assembly protein 1-like 4 (NAP1L4), serine/arginine-rich splicing factor 11 (SRSF11), and DNA topoisomerase 2-alpha (TOP2A), also were inhibited after treatment with CIGB-300, although the down-regulated phospho-residue was different in each AML cell line (**Table 1**, **Table S6**).

## 4. Discussion

After no considerable changes in AML therapy, growing knowledge in the molecular pathophysiology of this hematological disease is being reflected in therapeutics with the recent approval for the FDA of a number of novel agents [57]. Such advances have remarked the feasibility of molecular targeted approaches to treat AML, especially those that impinge on protein kinases with critical roles in the maintenance of malignant phenotype. Therefore, the exploration of kinase inhibitors with potential anti-neoplastic effect could significantly contribute to maintain the positive trend that we have witnessed in the last years [57]. Accordingly, here we showed that the clinical-grade CK2 inhibitor CIGB-300 impairs AML cells proliferation and viability, and provided mechanistic insights supporting this anti-leukemic effect.

Contrary to most kinases, which remain inactive until its activity is triggered by specific stimuli, CK2 is constitutively active and independent of second messengers or post-translational modifications [18]. Such distinctive feature makes possible for this kinase to keep oncogenic pathways constitutively activated, in a way that transformed cells develop an excessive “addiction” or reliance on its activity [58]. The abovementioned has fostered the evaluation of several CK2 pharmacologic inhibitors in pre-clinical cancer models, and more importantly, two of them have entered to clinical trials [25, 26]. Considering that CIGB-300 is one of the CK2 inhibitor undergoing clinical evaluation, we decided to explore its potential anti-leukemic effect against AML.

We found that the peptide inhibited AML cells proliferation in a dose-dependent manner, with mean IC_50_ of 27.9 μM. In contrast with solid tumor cells, which display an heterogenous sensitivity profile, leukemia cells are more sensitive to CIGB-300 anti-proliferative effect. Indeed, in a panel containing human lung, cervix, prostate and colon cancer cell lines, the IC_50_ values for CIGB-300 ranged between 20 and 300 μM [32], while in chronic and acute T-cell lymphocytic leukemia (CLL and T-ALL) the estimated values were all below 40 μM [34, 44].

Once evaluated the impact of the peptide on proliferation of five AML cell lines, HL-60 and OCI-AML3 cells were selected for further experimentation. These cell lines derived from the most common AMLs (i.e. acute promyelocytic and acute myelomonocytic leukemia), together accounting for roughly two thirds of all AML cases [59]. Importantly, in line with compelling evidence supporting that CK2 inhibition impairs cancer cells viability [20], we demonstrated that treatment with CIGB-300 leads to apoptosis in AML cell lines and primary cells. Moreover, cell cycle analysis indicated that HL-60 cells accumulated in S phase once treated with CIGB-300, whereas no changes in cell cycle distribution were detected in OCI-AML3 cells.

Protein kinase CK2 is a key regulatory player in signaling pathways that control proliferation, apoptosis cell death, and cell cycle progression [60]. Accordingly, we found that CK2 inhibition with CIGB-300 modulates members of the MAPKs and CDKs family, and impairs the CK2-mediated phosphorylation of CK2 substrates with key roles in proliferation and cell cycle. For instance, the filament-forming cytoskeletal GTPase SEPTIN2 is necessary for normal chromosome segregation and spindle elongation during the progression through mitosis, and its phosphorylation at S218 by protein kinase CK2 is crucial for cancer cell proliferation [61, 62]. The MCM2 subunit of the replicative helicase complex (MCM complex), which is essential for once per cycle DNA replication and cell division [63], was also found modulated in AML cells phosphoproteomic profiles. Phosphorylation of MCM2 at S41 and S139 (both sites down-phosphorylated in AML cells treated with CIGB-300), promotes the ATPase activity of the MCM helicase complex and the replication of DNA [64].

Noteworthy, the death promoting transcriptional repressor BCLAF1 was found inhibited at S397 in both AML cells. BCLAF1 was originally described as a protein interacting with anti-apoptotic members of the BCL2 family, and more recently it has been linked to cancer cells proliferation, invasion and drug resistance [65, 66]. Nevertheless, validation of BCLAF1 S397 as phosphorylation site targeted by protein kinase CK2, and the biological implications of such post-translational modification is something that need further experimentation.

In other vein, previous studies with the ATP-competitive inhibitor CX-4945 showed that protein kinase CK2 could regulate p53 protein turnover and PI3K/AKT signaling in AML cells [43, 50]. In such studies, HL-60 cells (which are p53 nulls), displayed refractoriness toward the cytotoxic effect of CX-4945, in part owing to the absence of PI3K/AKT modulation following treatment with CX-4945 [43, 50]. In contrast, herein we evidenced that HL-60 cells are similarly sensitive to CIGB-300-induced cell death when compared to OCI-AML3, thus suggesting the existence of divergent mechanism of action for each CK2 inhibitor in AML cells. In spite of such differences, in this study and previous ones, modulation of MAPKs and CDKs appeared as a common down-stream consequence of CK2 inhibition with both CIGB-300 and CX-4945 [33, 50].

The impact of CIGB-300 on ribosome biogenesis and protein translation has been suggested [31, 33]. Interestingly, here we evidenced that CIGB-300 is able to interact with several proteins from the small and the large ribosome subunits, probably due to the presence CK2 consensus sequence or acidic stretches in their structures. Similarly, phosphoproteomic analysis revealed that components of eukaryotic translation initiation factors such as EIF3C, EIF4G1 and EIF5B, were down-phosphorylated in AML cells. EIF3C is needed for the initiation of protein synthesis, specifically targeting and initiating translation of a subset of mRNAs related to cell proliferation, apoptosis, differentiation and cell cycle [67]. Furthermore, the catalytic subunit of H/ACA small nucleolar ribonucleoprotein complex (DKC1), which is required for ribosome biogenesis [68], was also identified among down-phosphorylated proteins in AML cells. Although NPM1 S125 phosphopeptide was not identified in our phosphoproteomic analysis, we demonstrated by western blot that CIGB-300 impairs its CK2-mediated phosphorylation. Nucleolar protein NPM1 is a molecular chaperone involved in diverse cellular processes such as ribosome biogenesis, mRNA processing, chromatin remodeling, mitotic spindle assembly and protein chaperoning [69].

In view of NPM1 has been suggested as a major target for the peptide in solid tumor cells [31], we decided to investigate if a similar mechanism could account in AML cells. Our results indicated that despite the clear interaction of this protein with the peptide and the subsequent inhibition of its CK2-mediated phosphorylation, those events do not appear to have a critical role in the anti-leukemic effect of CIGB-300. In fact, we achieved almost 85% knock-down of NPM1 protein with no significant changes in cell viability or sensibility to CIGB-300 anti-proliferative and cytotoxic effect in AML cells.

Accordingly, pull-down experiments and quantitative phosphoproteomics analysis have recently pointed out that the CIGB-300 inhibitory mechanism could be more complex than originally thought [33, 34]. In line with such results, here we evidenced that with the exception of NPM1 and NCL, which have been ruled out as major targets for the peptide [31, 34], none of the other CIGB-300-interacting CK2 substrates were identified as down-phosphorylated in AML cells. Instead, we showed that in AML cells the peptide is able to interact with CK2α catalytic subunit and regulate part of the CK2-dependent phosphoproteome, thus pointing at direct enzyme blockade as a putative inhibitory mechanism operating in AML. Whether this novel mechanism is universal and runs in parallel to substrate binding, or its relative contribution relies on the neoplastic cellular background is something that remains to be fully elucidated.

## 5. Conclusions

Here we explored for the first time the anti-leukemic effect of the clinical-grade synthetic-peptide CIGB-300 in AML. Importantly, we found that the peptide decreased the proliferation of AML cells in a dose-dependent manner, effect that could be explained by the induction of apoptosis and the impairment of cell cycle progression. To understand the antileukemic effect of CIGB-300, we further examined its internalization and subcellular localization. In agreement with the onset of cellular effect attained in AML cells, the peptide was rapidly internalized and distributed between the cytoplasm and the nucleus, with preferential accumulation in the nucleolus of HL-60 cells. Considering that NPM1 has been previously described as a major target for the peptide in solid tumor cells, we down-regulated this protein to clarify its relevance for the mechanism of action of CIGB-300. Consistent with NPM1 knock-down, the survey of CIGB-300 interactome and quantitative phosphoproteomic analysis, corroborated that CIGB-300 effect in AML could also be mediated by direct impairment of protein kinase CK2 enzymatic activity, in addition to binding to acidic phosphoacceptor in the substrates. Remarkably, a repertoire of CK2 substrates differentially modulated by CIGB-300 was identified, providing a molecular basis for the anti-neoplastic effect of the peptide against AML. In summary, our results not only revealed a number of mechanistic insights related to CIGB-300 anti-neoplastic effect in AML cells, but also highlighted the feasibility of protein kinase CK2 pharmacologic inhibition for AML targeted therapy.

## Author Contributions

Conceptualization, S.E.P., Y.P., D.V.-B and L.J.G.; methodology, V.B., Y.R., E.C., D.A., J.R.W. and K.Z.; formal analysis, M.R., A.R.-U. and V.B.; investigation, M.R., G.V.P., A.C.R. and Y.C.; writing—original draft preparation, M.R.; writing—review and editing, S.E.P., Y.P., D.V.-B and L.J.G.; supervision, S.E.P., Y.P. and Y.K.; project administration, S.E.P; funding acquisition, J.R.W., V.B., L.J.G., S.E.P. and Y.K. All authors have read and agreed to the published version of the manuscript.

## Institutional Review Board Statement

The study was conducted according to the guidelines of the Declaration of Helsinki, and approved by the Institutional Review Board of the Center for Genetic Engineering and Biotechnology (IG/CIGB-300I/LAR/1101, Havana, Cuba).

## Informed Consent Statement

Informed consent was obtained from all subjects involved in the study.

---

**Figure S1.**
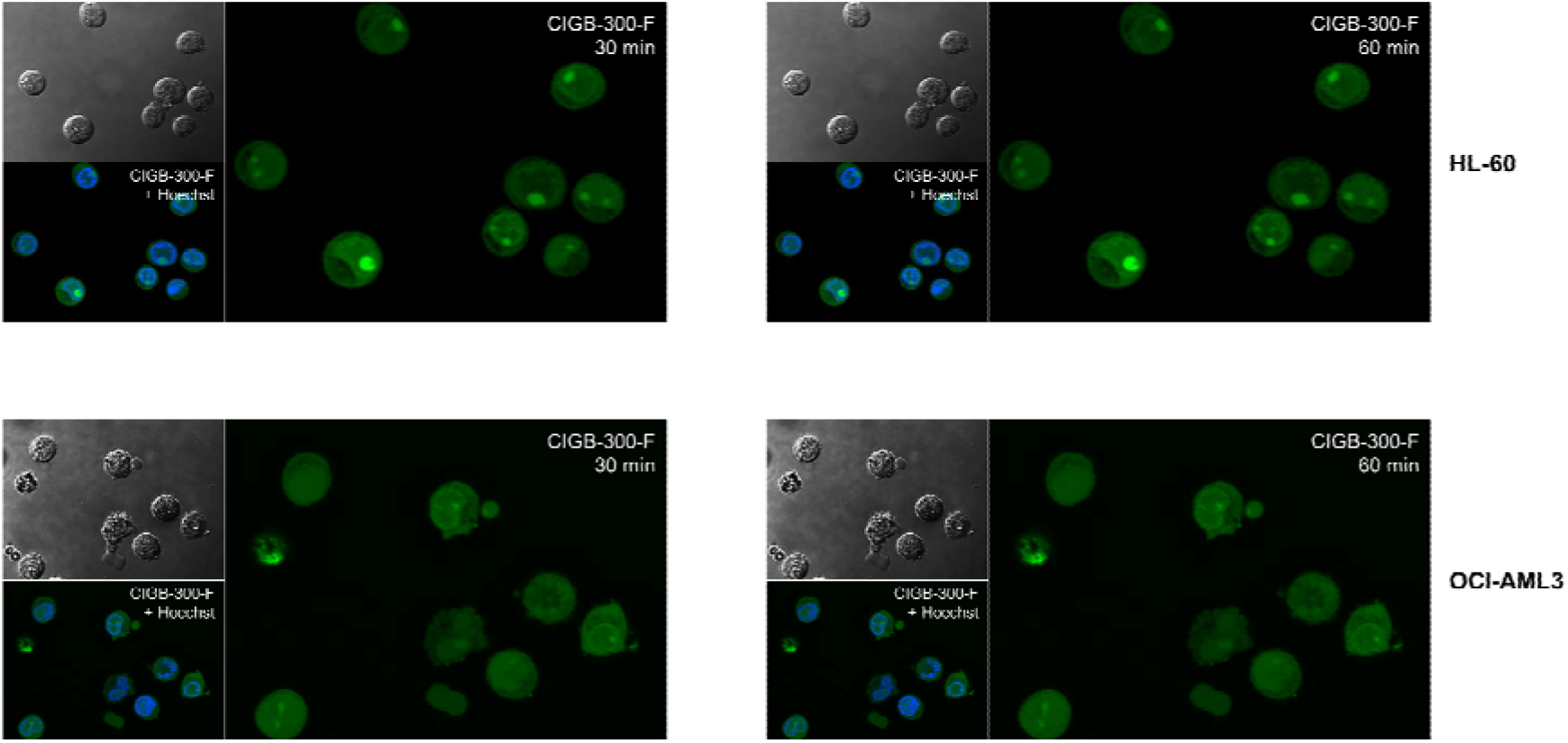
Subcellular distribution of CIGB-300 in AML cells after 30 and 60 min of incubation with 30 μM of CIGB-300-F. A total of 5 optical fields were examined for each experimental condition in confocal microscopy experiment.

**Figure S2.**
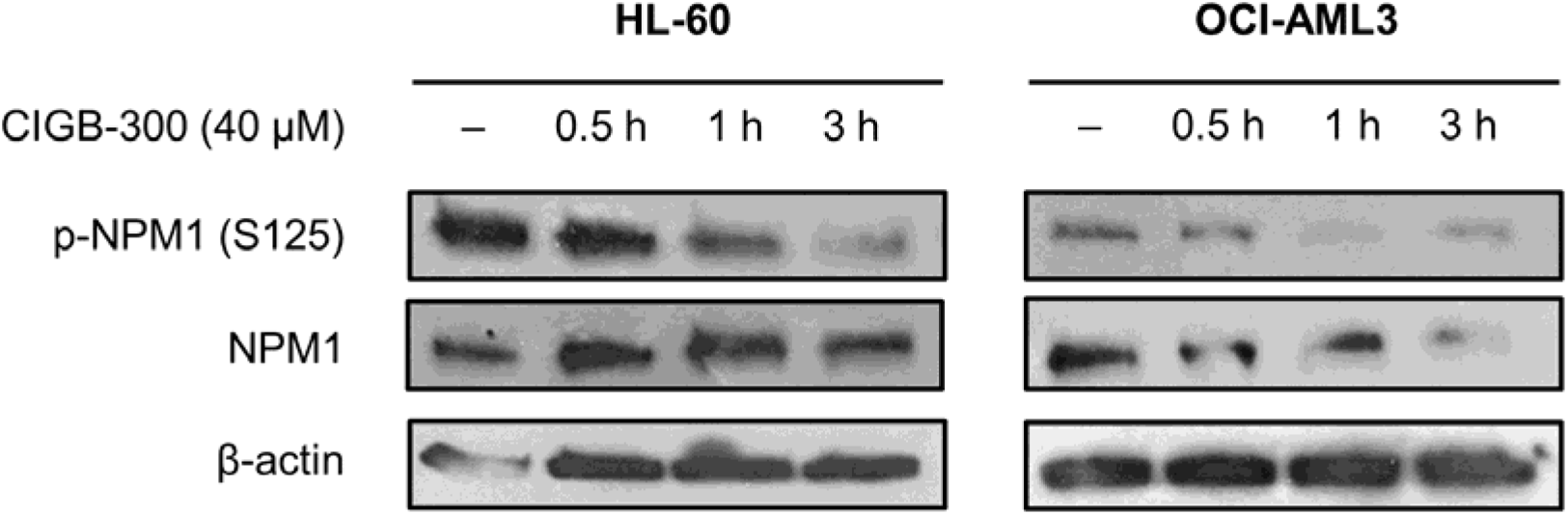
CIGB-300 peptide impairs CK2-mediated phosphorylation of NPM1 in AML cells. Cells treated with 40 μM of CIGB-300 during the indicated times were analyzed by western blot using phospho-specific and total NPM1 antibodies.

**Figure S3.**
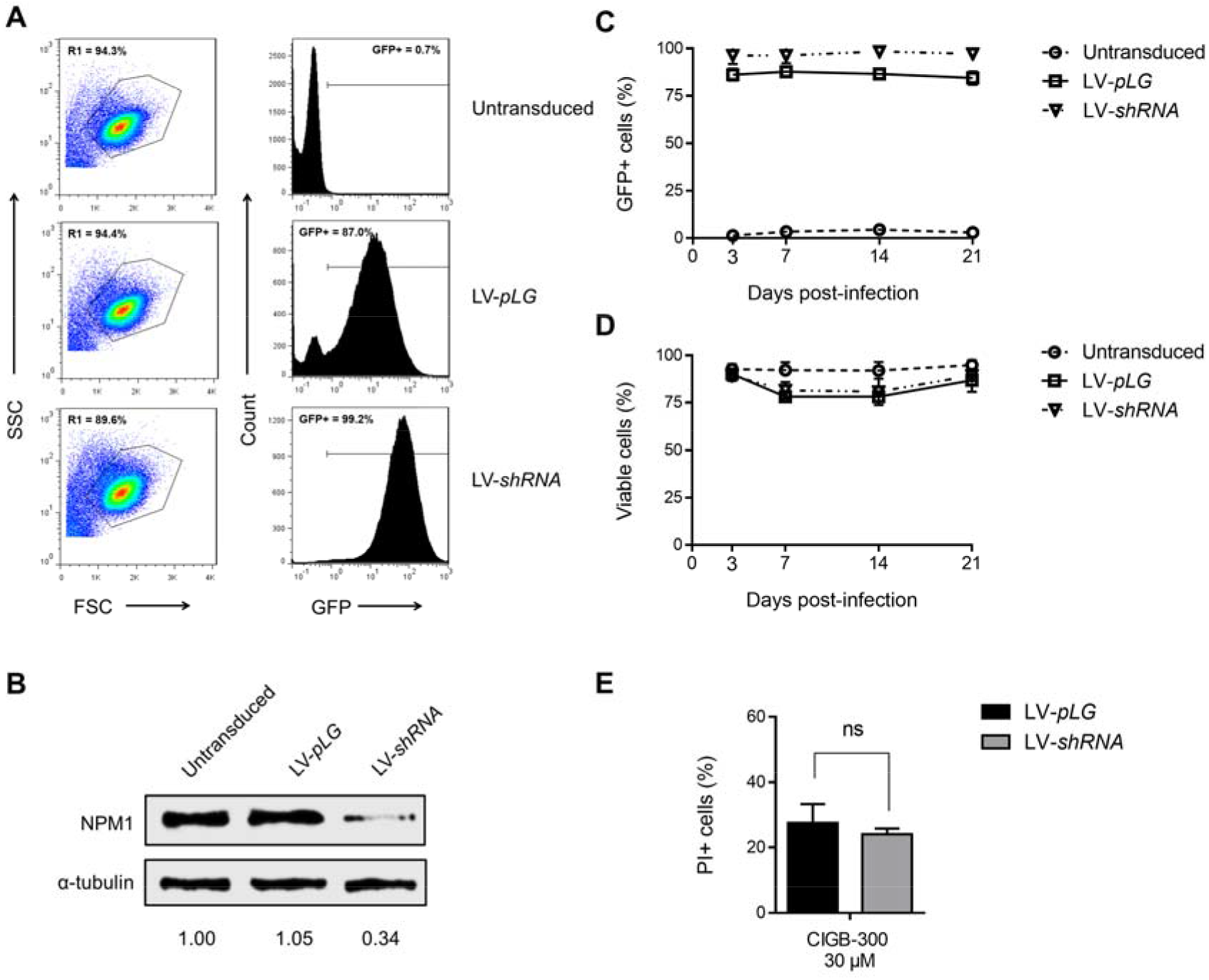
Lentiviral vector-mediated knock-down of NPM1 in THP-1 cells.: (**a**) Flow cytometry analysis of GFP expression in transduced cells at day 7 post-infection. Percentage of GFP+ cells was determined on live-cell population (R1 gate in dot plots); (**b**) Immunoblots from HL-60 transduced cells showing NPM1 protein knock-down in LV-*shRNA* infected cells; (**c**) GFP expression levels and (**d**) viability of transduced THP-1 cells were followed by flow cytometry during three weeks post-infection; (**e**) Sensibility of transduced cells toward the cytotoxic effect of CIGB-300 was evaluated by PI staining. Cells were incubated with 20 or 40 μM of CIGB-300 for 5 h, stained with PI solution and then analyzed by flow cytometry. Results from (**c**), (**d**) and (**e**) are shown as mean ± SD, n = 3. (ns) not significant.

**Table S1.** Characteristics of AML patients.

| Patient | Sex | Age | AML FAB subtype | Bone marrow blast |
| --- | --- | --- | --- | --- |
| 1 | Female | 76 years | FAB-M4 | 50 % |
| 2 | Male | 54 years | FAB-M2 | 80 % |
| 3 | Male | 59 years | FAB-M2 | 50 % |
| 4 | Female | 73 years | FAB-M5 | 90 % |
| 5 | Male | 47 years | FAB-M2 | 60 % |

**Table S2.** CIGB-300 interaction profile in AML cells.

**Table S3.** Phosphoproteome identified in AML cells.

**Table S4.** Phosphopeptides differentially modulated in AML cells treated with CIGB-300.

**Table S5.** Kinases related to differentially modulated phosphopeptides in AML cells treated with CIGB-300.

**Table S6.** Candidate CK2 substrates differentially modulated in AML cells treated with CIGB-300.

